# Joint estimation of selection intensity and mutation rate under balancing selection with applications to HLA

**DOI:** 10.1101/2021.11.18.469194

**Authors:** Montgomery Slatkin

## Abstract

A composite likelihood method is introduced for jointly estimating the intensity of selection and the rate of mutation, both scaled by the effective population size, when there is balancing selection at a single multi-allelic locus in an isolated population at demographic equilibrium. The performance of the method is tested using simulated data. Average estimated mutation rates and selection intensities are close to the true values but there is considerable variation about the averages. Allowing for both population growth and population subdivision do not result in qualitative differences but the estimated mutation rates and selection intensities do not in general reflect the current effective population size. The method is applied to three class I (HLA-A, HLA-B and HLA-C) and two class II loci (HLA-DRB1 and HLA-DQA1) in the 1000 Genomes populations. Allowing for asymmetric balancing selection has only a slight effect on the results from the symmetric model. Mutations that restore symmetry of the selection model are preferentially retained because of the tendency of natural selection to maximize average fitness. However, slight differences in selective effects result in much longer persistence time of some alleles. Trans-species polymorphism (TSP), which is characteristic of MHC in vertebrates, is more likely when there are small differences in allelic fitness than when complete symmetry is assumed. Therefore, variation in allelic fitness expands the range of parameter values consistent with observations of TSP.

---

The major-histocompatibility complex (MHC) in humans is called the human leucocyte antigen (HLA) system. The HLA system is found on the short arm of chromosome 6. Many HLA genes are involved in the adaptive immune response. Two groups of HLA loci have been intensively studied, class I and class II, which are similar but differ somewhat in function. They both code for glycoproteins that bind with peptides and present them at the cell surface to T cells and natural killer cells. They are key components in the body’s ability to defend itself against pathogens. They are important clinically because many HLA alleles are associated with higher risk of auto-immune and other heritable diseases and with susceptibility and resistance to various pathogens. (Trowsdale and Knight 2013) For example, the B27 allele at HLA-B is associated with a higher risk of ankylosing spondylitis (Evans *et al*. 2011) and alleles at the HLA-C locus are associated with different rates of HIV disease progression. (Kulpa and Collins 2011). HLA loci are also critically important to the success of organ transplants.

Class I and class II HLA loci are unusually diverse, with dozens and even hundreds of alleles at each locus distinguishable both by serotyping and DNA sequencing. (Radwan *et al*. 2020) Most of the variation among alleles is concentrated in the peptide binding regions, the second and third exons of the class I loci and the second exon of the class II loci. There is clear evidence of trans-species polymorphism (TSP): some alleles at both class I and class II loci in humans are more similar to alleles in chimpanzees than they are to other alleles at the same loci in humans. (Klein *et al*. 2007; Radwan *et al*. 2020) Both of these features have long been taken as evidence of strong balancing selection, meaning that on average individual heterozygous at HLA class I and II loci have a higher fitness than homozygous individuals.

Two types of balancing selection have been commonly invoked, heterozygote advantage and rare allele advantage (also called negative frequency-dependent selection). Both heterozygote advantage and rare allele advantage are assumed to result from the role class I and II loci play in the immune system. In general, alleles differ in their peptide binding regions and hence can present different antigens for inspection by T cells. Consequently, an individual that can present more kinds of antigens efficiently because it is more heterozygous will likely be able to mount an effective immune response to a larger variety of pathogens. The difference between the two hypotheses is, from a population genetics perspective, not very important. In the heterozygote advantage model, heterozygosity at each locus is itself favored by selection because it provides defense against a pathogen pool that is regarded unchanging. In the rare allele advantage model, an allele in low frequency is favored because pathogens are not yet well adapted to it. As an allele increases in frequency, pathogens adapt and the allele’s contribution to fitness decreases. In the rare allele advantage model, the HLA loci and pathogens coevolve in a way that results in higher average fitness of heterozygous individuals. In the heterozygote advantage model, coevolution plays no role. The reason the difference between the models is not important for population genetic analysis is that Takahata and Nei (1990) showed the equations that govern the change in allele frequency are the same in the two models when the allelic fitness in the rare allele advantage model is a linearly decreasing function of frequency. Spurgin and Richardson (2010) provided further support for Takahata and Nei’s conclusion.

There is extensive theory of balancing selection. I will review here only developments directly related to this paper. Kimura and Crow (1964) introduced the infinite-alleles mutation model, in which each mutation at a locus creates an allele new to the population. They also introduced the symmetric model of heterozygote advantage in which every homozygous individual has a fitness of 1-*s* relative to every heterozygous individual. Their model assumed a population of constant effective size, *N*. They showed that, for large *N*, the model’s behavior depends on only two parameters denoted here by, *θ* = 4*N*μ, where μ is the mutation rate, and *S* = 2*Ns*. I will call this model the symmetric overdominance model. It will serve as a baseline model with which other models will be compared.

Ewens (1972) showed that, for the infinite-alleles model with *s*=0, the number of alleles, *k*, in a sample is a sufficient statistic for estimating *θ*. Furthermore, he derived the sampling distribution of neutral alleles given *k* and the sample size. Watterson (1977) extended Ewens’s theory and derived the sampling distribution for overdominant alleles (*s*>0). Watterson’s sampling distribution is calculable only for small *s*, however. Watterson also showed that the computed homozygosity 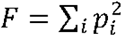, where *p_i_* is the frequency of the ith allele, is a powerful statistic for testing neutrality against the alternative of symmetric overdominance. The resulting test of neutrality is called the Ewens-Watterson test, which has been extensively applied to HLA data. (Solberg *et al*. 2008). Samples of class I and II HLA loci in many populations lead to rejection of neutrality, reinforcing the idea that diversity at HLA loci is maintained by balancing selection.

Takahata (1990) developed an analytic approximation for the symmetric overdominance model valid if *θ* « *S*. Takahata showed that under this assumption, gene genealogies in the symmetric overdominance model are similar to neutral gene genealogies but with a much larger effective population size. Takahata argued that for biologically reasonable parameter values, the symmetric overdominance model predicted large numbers of alleles and TSP.

Satta et al. (1994) used Takahata’s (1990) results to develop a method for estimating the intensity of selection in the symmetric model. Satta et al. (1994) estimated the mutation rate in the peptide binding region of class I and class II loci to be 1.7 × 10^-6^ and 1.5 × 10^-7^ respectively per generation. They then used Takahata’s (1990) analytic approximation combined with estimates of the non-synonymous substitution rate to estimate the selection intensity (*s*) of the major class I and class II loci to be between 0.0007 (for HLA-DPB1) and 0.042 (for HLA-B). Yasukochi and Satta (2013) revisited the problem using more extensive sequence data and confirmed the previous results. Yasukochi and Satta (2013) (Table 2) estimated *s* for HLA-A (0.0225), HLA-B (0.044) and HLA-DRB1 (0.0194) with smaller values for other loci. Assuming *N*=100,000, they concluded that *S* for these three loci were very large, (4500, 8825 and 3890 respectively). Yasukochi and Satta (2013) estimated the scaled mutation rates to be much smaller than 1, with θ between 0.04 and 0.92.

In this paper, I introduce a method for jointly estimating *θ* and *S* from the allele frequency spectrum under the assumption of symmetric overdominance in an equilibrium population. I then examine by simulation the dependence of the estimated parameter values when the assumptions of the symmetric model are relaxed in various ways. I will show that small variations in heterozygous fitness can make the long persistence of some alleles much more likely, which implies that TSP is facilitated by subtle variation in allelic contributions to fitness.

## Allele frequency spectrum

The basis for the method presented here is the allele frequency spectrum for the model of symmetric balancing selection. Muirhead and Wakeley (2009) derived the frequency spectrum for a population of 2*N* haploid individuals under frequency-dependent selection. They extended the Moran model to allow for multiple alleles and linear dependence of the death rate on allele frequency (their selection-at-death model). In this model, the probability that an allele decreases by one copy is proportional to 1 – *sx*, where *s* is the selection intensity and *x* is the allele frequency. They relied on Takahata and Nei’s (1990) result that the linear dependence of fitness on allele frequency is formally equivalent to symmetric balancing selection. Muirhead and Wakeley (2009) derived the expectation of the allele frequency spectrum, which is the number of alleles in each frequency class, given the scaled mutation rate *θ* = 2*Nu* and the scaled selection intensity *S* = *Ns*. Because the effective size of a Moran model is smaller by a factor of 2 than the effective size of a Wright-Fisher model with the same number of individuals (Muirhead and Wakeley 2009), their results are applicable to a diploid population containing N individuals.

Muirhead and Wakeley (2009) showed that the spectrum satisfies the system of equations

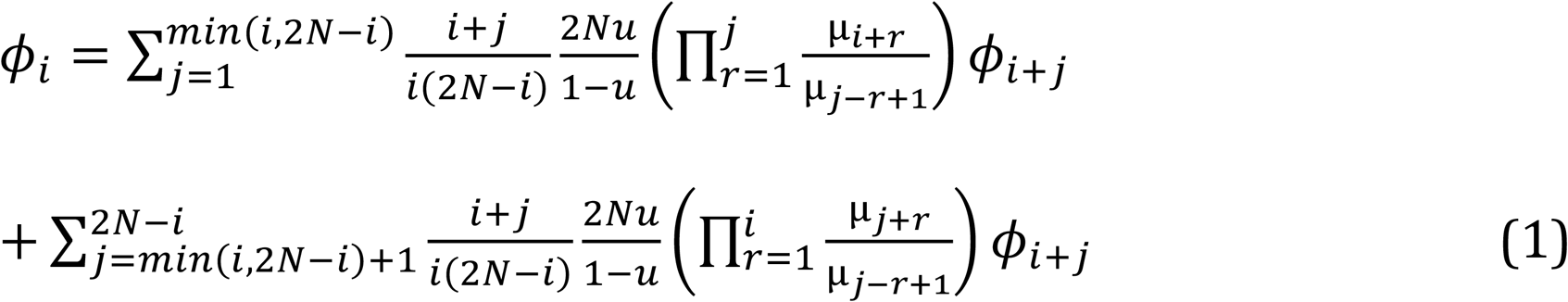

where *ϕ_i_* (*i* = 1,2,…,2*N*) is the expected number of alleles present in *i* copies and μ_j_ = 1 + *sj*/(2*N*). These equations can be solved iteratively by choosing an arbitrary value of *ϕ*_2*N*_ and then computing *ϕ_i_* for successively smaller values of *i* until *i* = 1 is reached. For reasons that will be clear later, the normalized spectrum of polymorphic alleles will be needed, so the value of *ϕ*_2*N*_ is determined by imposing the condition 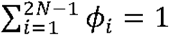.

The *ϕ_i_* represent the frequency spectrum in the whole population. The spectrum in a sample of *n* individuals is obtained by assuming that 2*n* copies are randomly chosen with replacement from the population. Therefore, the probability that there are *j* copies in the sample given *i* copies in the population is the binomial distribution

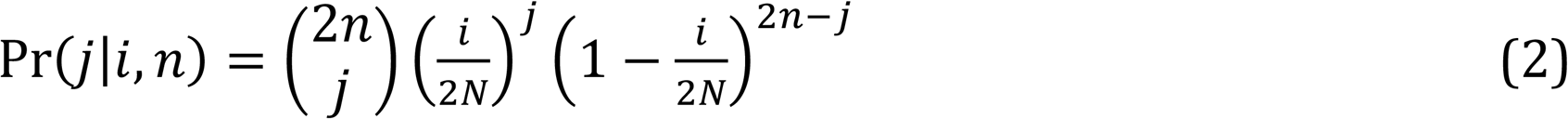

By summing over *i*,

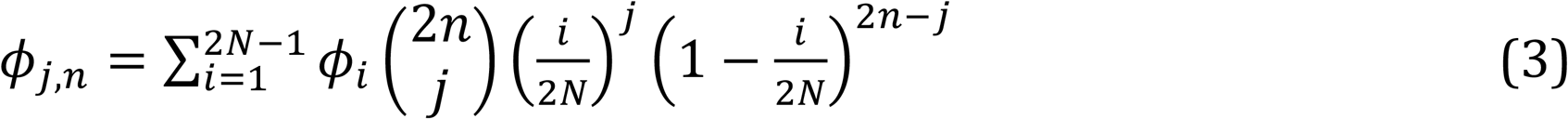

for *j* = 1,…,2*n* – 1. The subscript *n* is added to indicate the sample size. The term for *i*=2*N* is omitted in the sum because fixed loci are ignored. Because of the normalization condition for 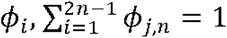 for any *n*.

## Composite likelihood method

The data for a single locus is the allele frequency spectrum in a sample of *n* individuals, *D* = {*i*_1_, *i*_2_,…} where *i_j_* is the number of alleles found in *j* copies (∑*_j_ i_j_* = *k*, ∑*_j_ ji_j_* = 2*n*). The probability of *D* can be approximated by assuming the *i_j_* are a random draw from a multinomial distribution with probability vector *ϕ_j,n_* and sample size 2*n*:

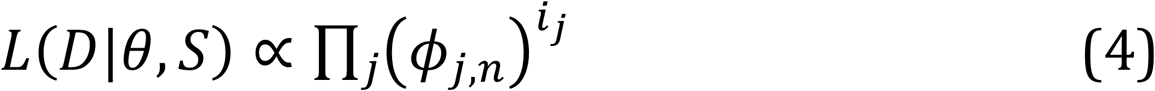

The result is a composite likelihood because it assumes independence of alleles when in fact they are not independent; their frequencies have to sum to 1. The *ϕ_j,n_* depend on *S* and *θ*. The maximum likelihood values of the two parameters, *Ŝ* and 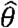, can then be found numerically. In the results presented below, a grid search with spacing 1 for *θ* (with a range 1-40) and 20 for *S* (with a range 0 to 1000) was used.

## Simulation tests

The first test of this method assumes the symmetric equilibrium model. A Wright-Fisher model with *N*=10,000 diploid individuals was subject to balancing selection. Every heterozygote had a relative fitness of 1 and every homozyote had a relative fitness of 1-*s*. Mutations occurred with probability *μ* per generation. Each mutant allele was new to the population. The population was initially fixed for a single allele. After a burn-in period of 100,000 generations, each replicate simulation continued for 1,000,000 generations with samples taken with replacement every 10,000 generations. Ten replicates were run for each set of parameter value. Averages were taken over the 1010 samples.

For each sample, *Ŝ* and 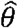 were obtained by using a grid search described above. In addition to estimating *S* and *θ, k* and *F* were computed. From *k, F* and *n*, the one-tailed Ewens-Watterson test (Watterson 1977) yielded the probability *P* that the observed value of *F* is smaller than a random value obtained by simulations with *s*=0.

Figure 1 shows the averages of 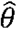 and *Ŝ* for a range of values of *S* and *θ*. The pattern in the results shown in Figure 1 are representative of other values of *θ*. Averages of 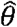 are close to the true values for a wide range of selection intensities, except when *θ* = 2 or smaller and *S* is relatively small. The average of *Ŝ* is close to the true value except for the largest value of *S*. The downward bias for large *S* results from the fact that the upper limit of the search interval for *Ŝ* was 1000. Figure 2 shows that the range of variation in 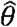 is not large both for relatively weak (*S* = 50) and relatively strong (*S* = 500) selection. In contrast, Figure 3 shows that the range of variation in *Ŝ* across samples is quite large. The extensive variation in *Ŝ* reflects the stochastic variation in allelic configurations across samples even when selection is strong. We can conclude from these results that if the equilibrium symmetric model is valid, *θ* can be estimated with some accuracy but individual estimates of *S* are probably not accurate.

**Figure 1.**
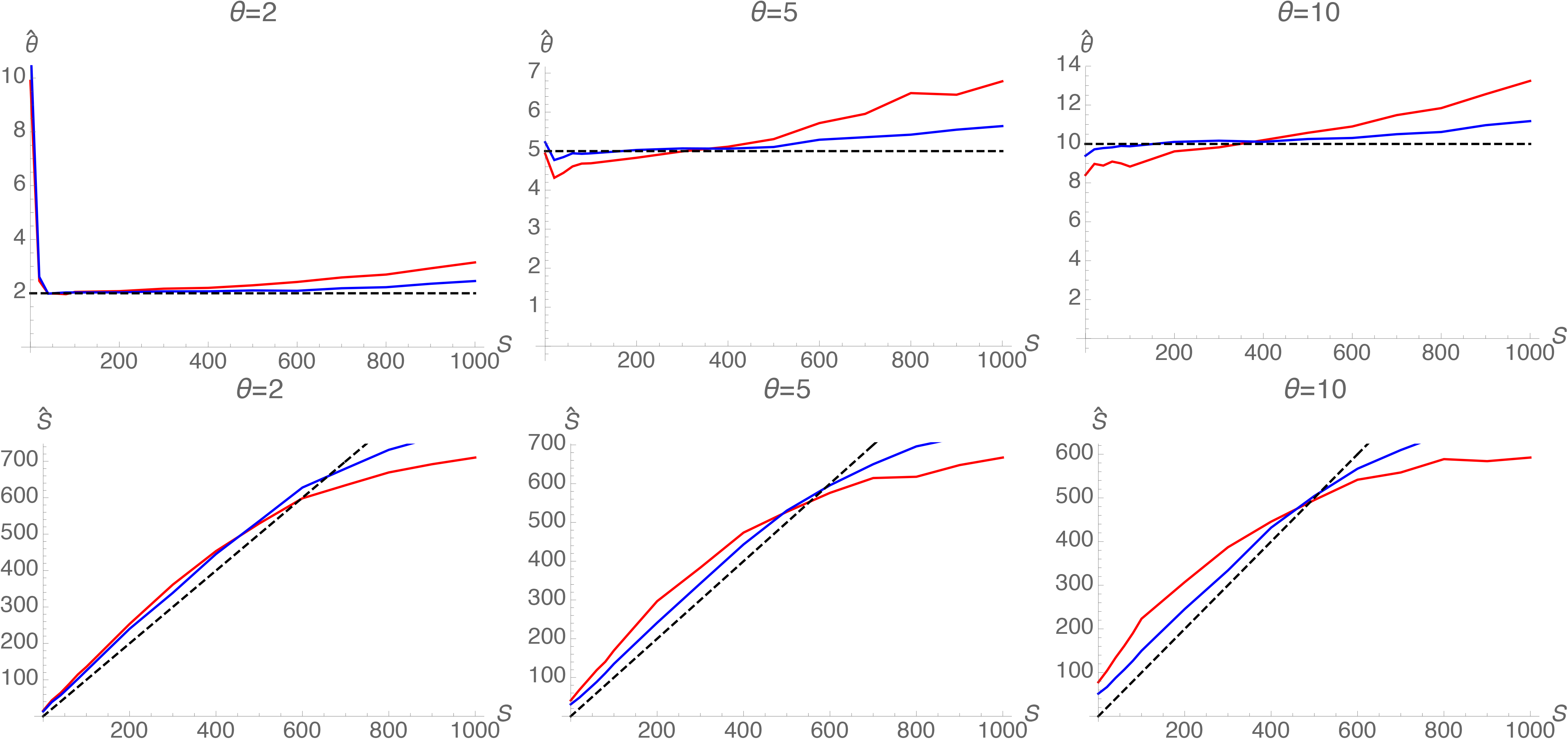
Average estimates of *θ* and *S* in a stable population of 10,000 individuals. In each of 10 replicates, there was a 100,000 generation burn-in period. Then 101 samples were drawn every 10,000 generations beginning with the end of the burn-in. The averages shown are over 1010 samples of 50 (red) and 500 (blue) individuals. The dashed horizontal lines show the true values of θ in each case. The dashed diagonal lines show the true values of *S*.

**Figure 2.**
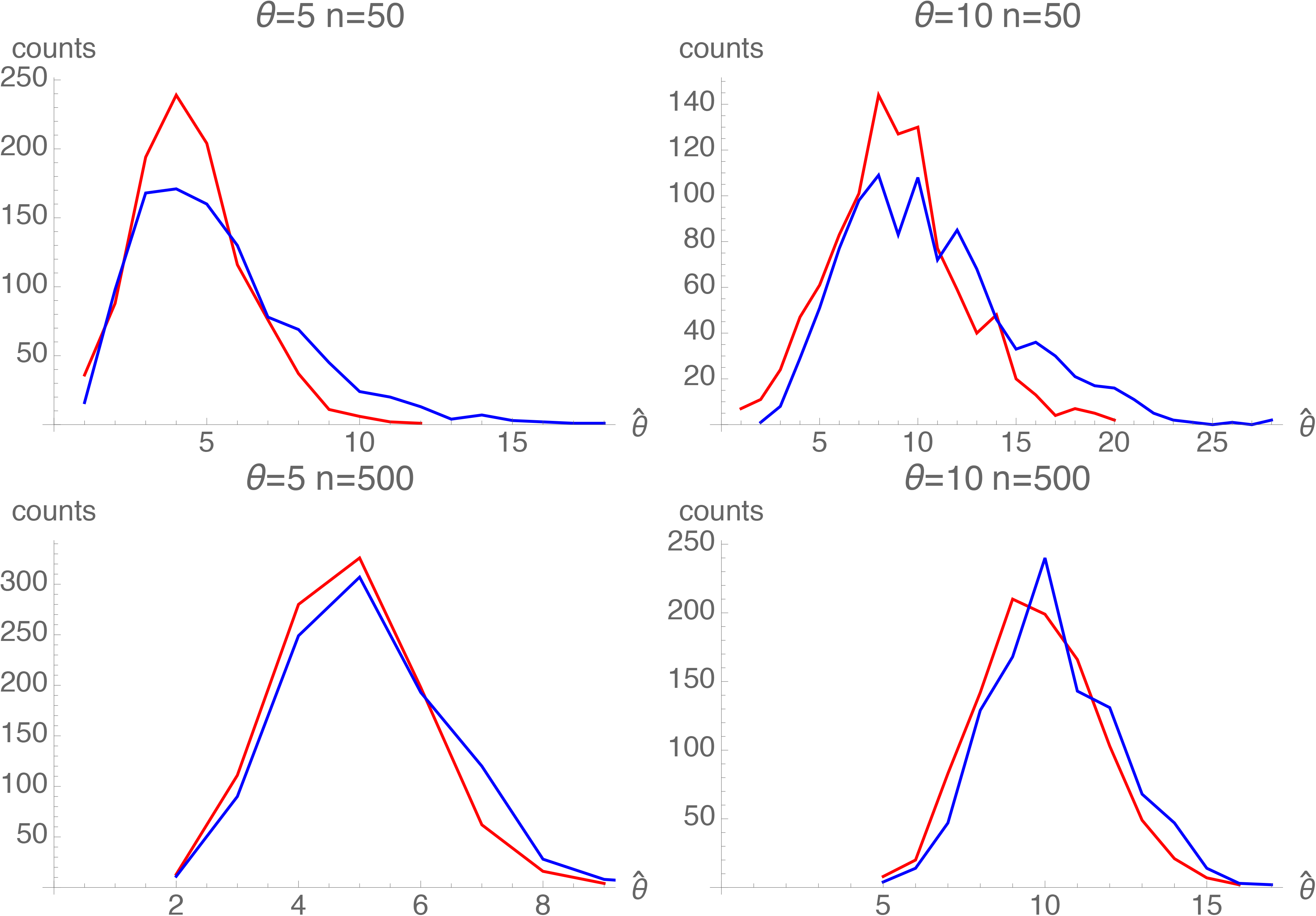
Distribution of 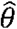 in samples from a population of size 10,000. The simulations are the same as those presented in Figure 1. The distribution of the 1010 sample values are shown for *S*=50 (red) and *S*=500 (blue).

**Figure 3.**
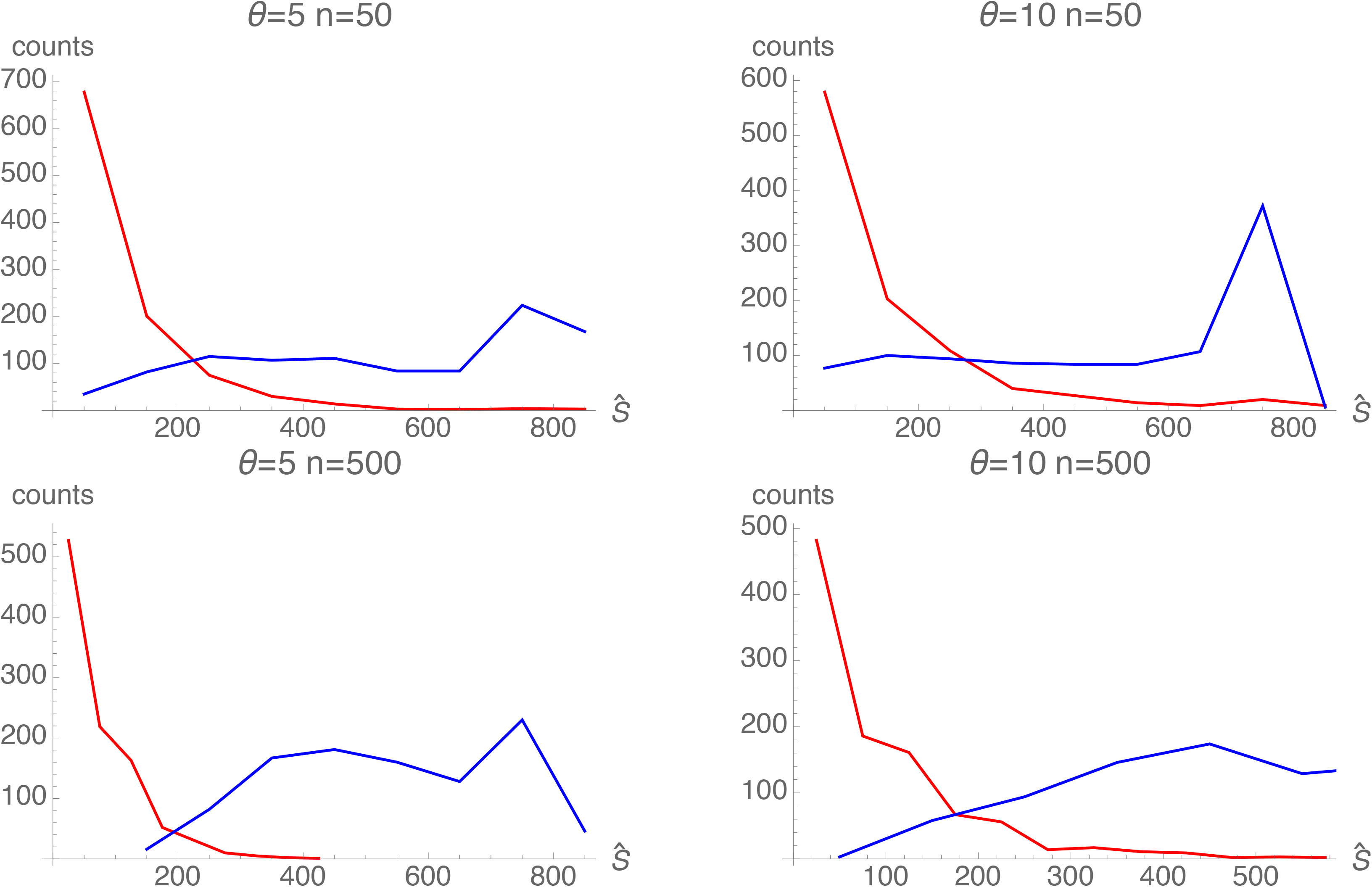
Distribution of *Ŝ* in from a population of size 10,000 The simulation results are the same as those presented in Figure 1. Distributions of the 1010 sample values are shown for *S*=50 (red) and *S*=500 (blue).

Figure 4 shows that the power of the Ewens-Watterson test (Watterson 1977) to reject neutrality decreases as *θ* increases. A comparison of the results in Figures 3 and 4 indicates that when values of *Ŝ* exceed 100, the Ewens-Watterson test has some power to reject neutrality. But unless θ is small and both *S* and *n* are large, the test does not in general have much statistical power. That conclusion is consistent with the results of Solberg et al. (2008) and others. For HLA loci in many populations, neutrality is rejected in some but not all cases.

**Figure 4.**
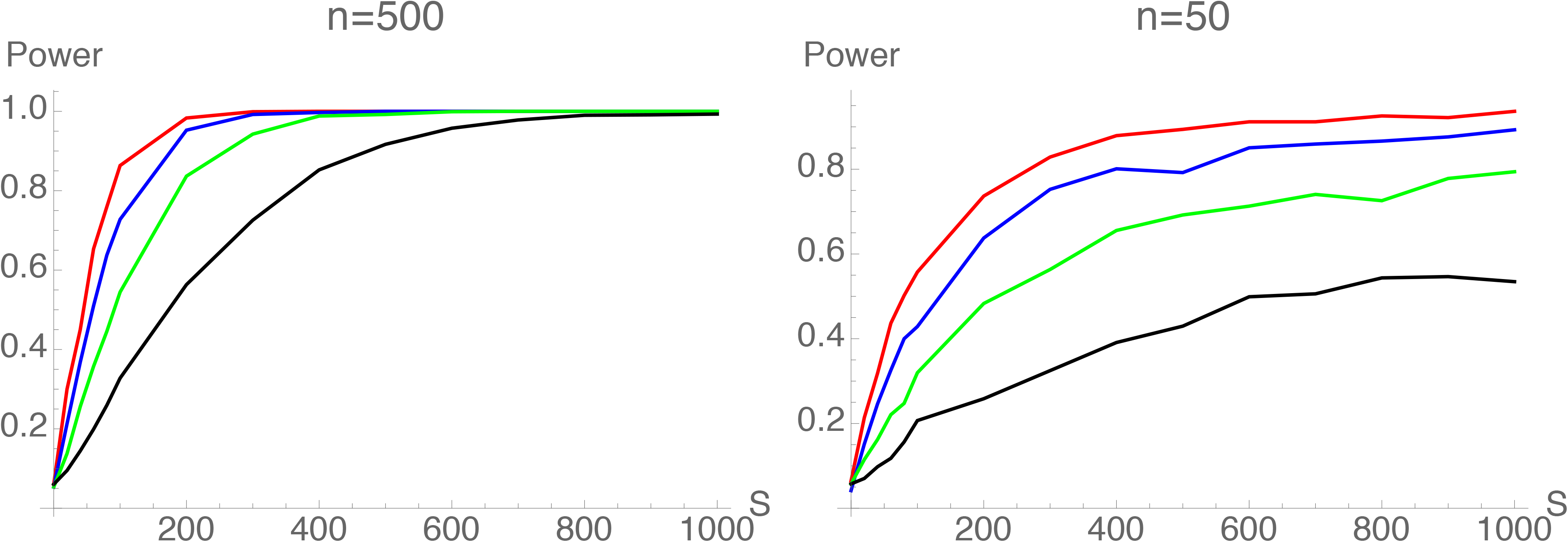
Power of Ewens-Watterson test to reject the hypothesis of neutrality at the 5% level (one-sided test) in the simulated data sets for a stable population with samples of *n*=50 and 500 individuals. The simulations are the same as those presented in Figures 1-3. Both parts show results for *θ* =2 (red), 3 (blue), 5 (green) and 10 (black).

## Application of the composite likelihood method

The composite likelihood method was applied to the data of Gourraud et al (2014). They used Sanger sequencing to determine the HLA genotypes at five loci (HLA-A, HLA-B, HLA-C, HLA-DQB1, and HLA-DRB1) of 930 individuals in the 1000 Genomes panel. They distinguished alleles by both the serotype and the sequence of the peptide binding region. Thus, for example, HLA-A*0101 was treated as a different allele than HLA-A*0102; 01 and 02 represent different amino acid sequences of the peptide binding region the HLA-A*1 allele. The data were extracted from Supplementary Table 2 of the Gourraud et al. paper. Data were available for 14 populations, two from Africa (LWK, YRI), four from the Americas (ASW, CLM, MXL PUR), four from East Asia (CHB, CHD, CHS, JPT) and four from Europe (CEU, FIN, GBR, TSI).

Table 1 shows the result of applying the composite likelihood method to the Gourraud et al. data set. Also indicated are the samples for which the Ewens-Watterson test rejected neutrality at a 5% significance level. Estimates of *θ* for each locus are roughly consistent across populations. Estimates for HLA-B are generally the largest and estimates for HLA-DQB1 are generally the smallest. Estimates for HLA-A, HLA-C and HLA-DRB1 are intermediate.

**Table 1.** Analysis of the 1000 Genomes data from Gourraud et al. (2014). Each cell contains the sample size (in italics) 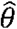 (in plain text), *Ŝ* (in bold) and the product *kF* (in bold italics). An asterisk indicates that the P value from the Ewens-Watterson is less than 0.05 (based on 1000 replicate simulations of the neutral distribution of *F*).

|  | Africa |  | America |  |  |  | East Asia |  |  |  | Europe |  |  |  |
| --- | --- | --- | --- | --- | --- | --- | --- | --- | --- | --- | --- | --- | --- | --- |
|  | LWK | YRI | ASW | CLM | MXL | PUR | CHB | CHD | CHS | JPT | CEU | FIN | GBR | TSI |
| A | *90* | *60* | *53* | *70* | *59* | *70* | *90* | *90* | *100* | *91* | *75* | *100* | *96* | *90* |
|  | 4 | 5 | 5 | 10 | 5 | 7 | 4 | 5 | 4 | 3 | 4 | 3 | 4 | 7 |
|  | **260\*** | **140** | **260\*** | **20** | **100** | **80** | **20** | **20** | **20** | **20** | **20** | **20** | **20** | **20** |
|  | **1.77** | **1.81** | **1.67** | **2.6** | **1.87** | **2.06** | **2.36** | **2.86** | **2.52** | **3.04** | **3.37** | **3.24** | **2.65** | **2.71** |
| B | *90* | *60* | *53* | *70* | *59* | *70* | *90* | *90* | *100* | *91* | *75* | *90* | *96* | *90* |
|  | 6 | 9 | 18 | 10 | 20 | 13 | 11 | 13 | 10 | 4 | 10 | 4 | 5 | 8 |
|  | **220\*** | **80** | **20** | **700\*** | **500** | **260** | **140** | **40** | **20** | **340\*** | **20** | **140\*** | **260\*** | **300\*** |
|  | **1.89** | **2.02** | **2.24** | **1.66** | **1.67** | **1.88** | **2.11** | **2.40** | **3.23** | **1.71** | **2.53** | **1.91** | **1.87** | **1.89** |
| C | *90* | *60* | *53* | *70* | *59* | *70* | *90* | *90* | *100* | *91* | *75* | *90* | *96* | *90* |
|  | 4 | 4 | 6 | 3 | 4 | 5 | 2 | 6 | 4 | 2 | 3 | 1 | 2 | 4 |
|  | **80** | **20** | **40** | **140\*** | **100** | **100** | **160\*** | **20** | **20** | **120\*** | **120\*** | **220\*** | **240\*** | **120** |
|  | **2.01** | **2.15** | **2.20** | **1.77** | **1.92** | **1.96** | **1.77** | **2.55** | **2.41** | **1.67** | **1.82** | **1.58** | **1.62** | **1.99** |
| DRB1 | *90* | *60* | *53* | *70* | *59* | *70* | *90* | *90* | *100* | *91* | *75* | *100* | *96* | *90* |
|  | 3 | 4 | 4 | 9 | 6 | 11 | 7 | 7 | 6 | 4 | 7 | 4 | 4 | 6 |
|  | **140\*** | **260\*** | **780\*** | **420\*** | **360\*** | **120** | **140** | **20** | **40** | **160** | **20** | **60** | **100** | **140** |
|  | **1.82** | **1.64** | **1.44** | **1.75** | **1.68** | **2.02** | **2.04** | **2.65** | **2.40** | **1.89** | **2.73** | **2.19** | **2.04** | **1.99** |
| DQB1 | *90* | *60* | *53* | *70* | *59* | *70* | *90* | *90* | *100* | *91* | *75* | *100* | *96* | *90* |
|  | 2 | 1 | 2 | 2 | 2 | 2 | 1 | 2 | 2 | 2 | 2 | 1 | 1 | 2 |
|  | **20** | **80** | **60** | **80\*** | **40** | **60** | **180\*** | **40** | **60** | **120\*** | **40** | **140\*** | **120\*** | **80\*** |
|  | **2.29** | **1.69** | **1.69** | **1.67** | **1.84** | **1.88** | **1.61** | **1.88** | **1.86** | **1.79** | **1.82** | **1.40** | **1.56** | **1.84** |

**Table 2.** Numbers of alleles that had ages of 100,000 generations or longer in the Vhet and Vpli models. The total numbers of mutations are in the denominator. Alleles were counted only if they arose after the end of the 100,000 generation burn-in period and were lost before the end of the 1,000,000 generations after the end of the burn-in period.

| Model | $\theta$ | $u_i$ uniform on (0,1) | | $u_i = 1$ | |
| --- | --- | --- | --- | --- | --- |
| | | $S=500$ | $S=1000$ | $S=500$ | $S=1000$ |
| Vhet | 0.1 | 15/50,085 | 17/50,017 | 39/48,689 | 47/50,327 |
| Vhet | 0.5 | 31/249,832 | 40/249,838 | 2/249,924 | 0/250,774 |
| Vhet | 1 | 25/500,888 | 42/500,771 | 0/501,016 | 0/499,083 |
| Vhet | 3 | 17/1,497,757 | 33/1,501,435 | 0/1,498,825 | 0/1,499,579 |
| Vhet | 5 | 6/2,496,873 | 15/2,501,992 | 0/2,498,105 | 0/2,499,456 |
| | | $r_i$ uniform on (0,0.05) | | $r_i = 0.0$ | |
| | | $S=500$ | $S=1000$ | $S=500$ | $S=1000$ |
| Vpli | 0.1 | 6/55,219 | 13/54,957 | 26/55,089 | 47/54,486 |
| Vpli | 0.5 | 22/275,095 | 26/275,048 | 1/275,297 | 0/275,111 |
| Vpli | 1 | 21/550,670 | 41/550,464 | 0/550,459 | 0/551,508 |
| Vpli | 3 | 38/1,650,207 | 39/1,650,541 | 0/1,651,103 | 0/1,650,563 |
| Vpli | 5 | 35/2,751,780 | 36/2,751,080 | 0/2,750,732 | 0/2,747,799 |

Estimates of *S* are much more variable than estimates of *θ*, as would be expected from Figure 3. Estimates for HLA-B tend to be the largest. With the exception of the two African populations, estimates for HLA-A tend to be the smallest. Neutrality can be rejected in some populations for all loci. Because of the dependence of the power of the Ewens-Watterson test on the mutation rate (see Figure 3), neutrality is rejected for HLA-DQB1 only for large *Ŝ*.

## Deviations from the symmetric equilibrium model

The symmetric equilibrium model is relatively easy to analyze but it is not appropriate for human HLA data for several reasons. The demographic assumptions, that human populations have been of constant size for a long time and isolated from one another, are obviously false. Human populations have grown rapidly in the recent past and have experienced bottlenecks in size in the more distant past (Li and Durbin 2009; Spence *et al*. 2018). Human populations show a pattern of isolation-by-distance both within and among geographic regions (Auton *et al*. 2009). The symmetry of the selection model is also unrealistic. Complete equivalence of alleles that are part of such a complex biological process seems implausible a priori. Models of interactions with pathogens do not result in symmetry (Borghans *et al*. 2004; De boer *et al*. 2004; Siljestam and Rueffler 2019; Stefan *et al*. 2019). And empirical evidence of various kinds argues against symmetry (Bronson *et al*. 2013; Paterson *et al*. 1998; Hedrick 2002; Ilmonen *et al*. 2007; Stoffels and Spencer 2008; Radwan *et al*. 2020)

Here, I consider the effects of deviations from the symmetric equilibrium model, first relaxing the demographic assumptions and then relaxing the assumption of symmetric balancing selection.

### Population growth

I simulated a model of exponential population growth by having a burn-in period of 100,000 generations (as in the simulations of a stable population). Then exponential growth occurred for 10,000 generations with a specified doubling time. Five samples were drawn, at 0, 2500, 5000, 7500 and 10,000 generations after the end of the burn-in period. One hundred replicates for each set of parameter values were run.

Figure 5 shows some results. Values of 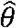 increase with the population size but not as rapidly as 4*N*(*t*)μ. The results are different for *Ŝ*. There is little systematic increase with time. The intuitive reason is that *Ŝ* depends mainly on alleles in relatively high frequency. Those alleles were already in high frequency at the onset of population growth and growth did not change their frequencies by much. In contrast, 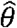 depends primarily on the numbers of low frequency alleles. Such alleles tend to be young and reflect the recent influx of mutations, which increases with the population size.

**Figure 5.**
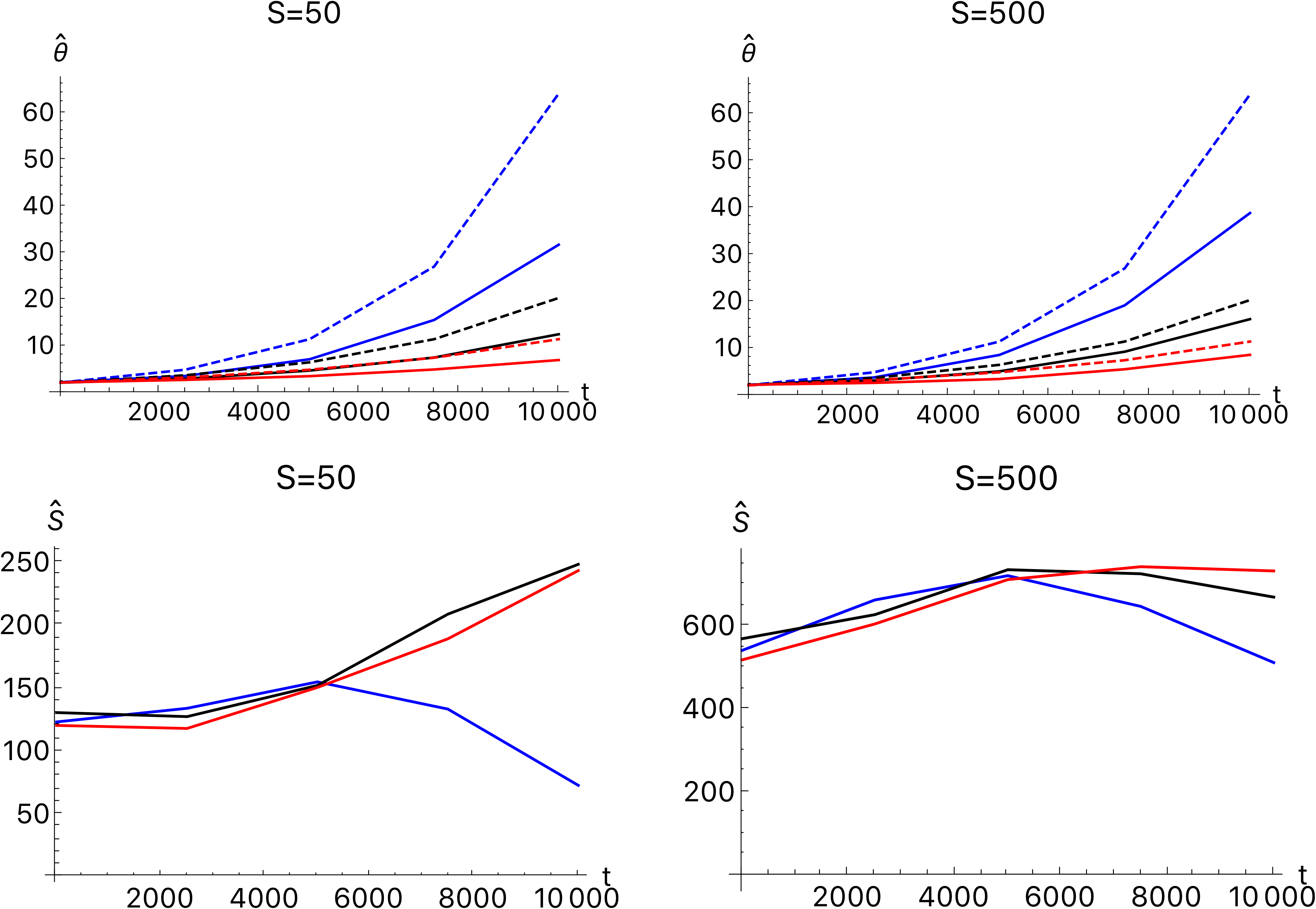
Estimates of θ and *S* for model of exponential population growth with the true value of *θ*=4*N*_0_μ=2 and three doubling times, 2000 (blue), 3000 (black) and 4000 (red) generations, where *N*_0_ = 10,000 is the population size during the 100,000 generation burn-in period. The dashed lines show 4*N*(*t*)μ for the three growth rates in colors corresponding to the solid lines.

### Population subdivision

As in the other simulations, there was a 100,000 generation burn-in period in which there was a single population of size 10,000. That population gave rise to an island model: five descendant populations, each of size *N*=10,000, exchanged migrants with one another each generation at a rate *m*/4. In the results, the migration rate is scaled by 4*N*, *M* = 4*Nm*. Samples of 500 individuals were drawn with replacement from each population every 10,000 generations and analyzed as if they were independent replicates. In addition, a sample of 500 individuals was drawn with replacement from all five populations together.

Some results are shown in Figure 6 for *M*=19, for which the expected value of Wright’s *F_ST_* is 1/(1 + *αM*), where *α* = *n*^2^/(*n* – 1)^2^ and *n* is the number of populations. (Crow and Aoki 1984). In this case, *n*=5 and the expectation of *F_ST_* is 0.032 which is a typical value for human populations within the same continent. (Elhaik 2012). Estimates of θ for each subpopulation are larger than the true value. The reason is that immigration introduces low frequency alleles that are equivalent to new mutations. Estimates of θ for the pooled sample are appropriately larger and of the same order of magnitude as the net mutation rate in the five populations together (10Aμ).

**Figure 6.**
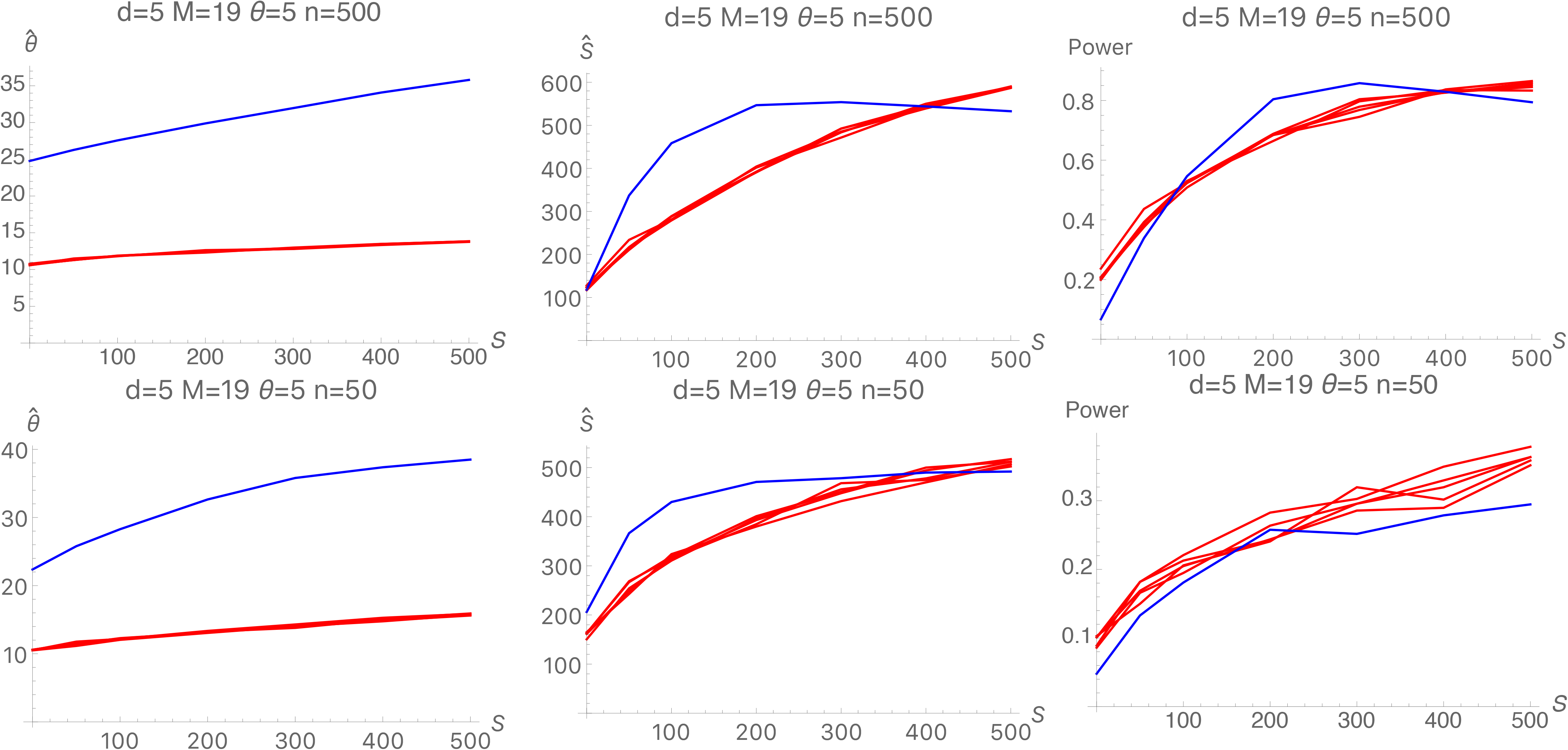
Estimates of *S* and θ along with statistical power to reject neutrality in an island model with *d*=5 subpopulations and a scaled migration rate of *M*=19. Results from analyzing each of the five populations separately are shown in red and the results from analyzing a sample from the five subpopulations mixed together are shown in blue.

For smaller values of *S, Ŝ* is biased upwards in each population and in the pooled population. Population subdivision tends to make the distribution of allele frequencies more even than in a single population, a pattern indicative of balancing selection that is reflected in larger *Ŝ*.

### Models of asymmetric balancing selection

To examine the effects of non-equivalence of HLA alleles, I simulated three models of asymmetric balancing selection. They do not exhaust the range of possibilities but they illustrate some general properties of such models.

The three models are (1) the variable homozygous fitness model (Vhom), (2) the variable heterozygous fitness model (Vhet), and (3) the variable pleiotropy (Vpli) model. In the Vhom model, each heterozygous individual has a relative fitness of 1 (as in the symmetric model) but each mutant allele is assigned a homozygous fitness of 1 – *s*(1 + *v_i_*) where *v_i_* is drawn from a uniform distribution on (0, *v_max_*). In the simulation results presented here, *v_max_* = 1. In the Vhet model, every homozygote has a relative fitness of 1 – *s* (as in the symmetric model) and every heterozygote has a relative fitness of 1 + *s*(*u_i_* + *u_j_*)/2, where *u_i_* is attached to the ith allele when it appears as a new mutation: *u_i_* is drawn from a uniform distribution on (0, *u_max_*). In the results presented here, *u_max_* = 1. In the Vpli model, symmetric balancing selection of intensity *s* is assumed. In addition, each mutation has a pleiotropic effect that reduces relative fitness by a factor 1 – *r_i_*, where *r_i_* is drawn from a uniform distribution on (0, *r_max_*). The pleiotropic effect is multiplicative: an *A_i_A_i_* individual has relative fitness (1 – *s*)(1 – *r_i_*)^2^ and an *A_i_A_j_* individual has relative fitness (1 – *r_i_*)(1 – *r_j_*). In the simulation results *r_max_* = 0.05.

It will be useful to see the dependence of the average fitness on the selection parameters in each model. Assume in a population that there are *k* alleles and the frequency of allele *A_i_* is *p_i_*. It is straightforward to show that if the genotypes are in their Hardy-Weinberg frequencies, the mean relative fitnesses for the three models are

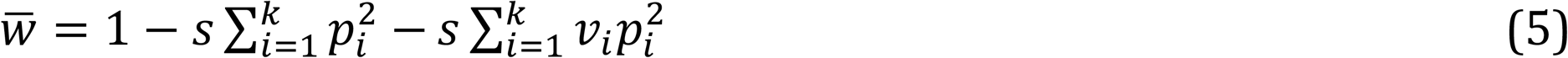

for the Vhom model,

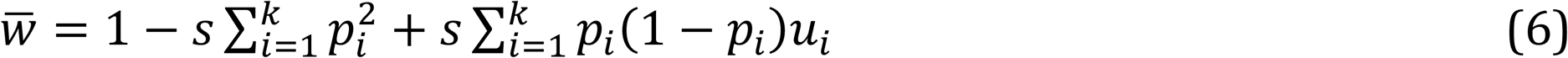

for the Vhet model, and

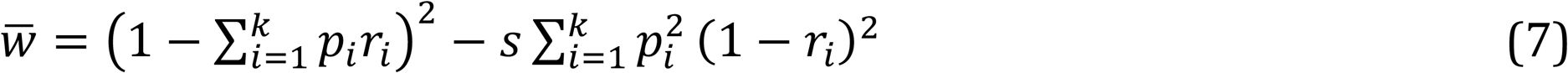

for the Vpli model. For a given set of allele frequencies, 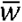 in Eq. (5) is a decreasing function of the *v_i_*. In Eq. (6), 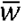 is an increasing function of the *u_i_* and in Eq. (7), 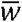 is a decreasing function of the *r_i_*.

Figure 7 shows that the results from simulations of the three models and a comparison with the comparable symmetric model. Results from the Vhom model (black curves) are very similar to those from the symmetric model (red curves). The Vhet model (blue curves) reduces *k* (the number of alleles) and 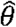, but has little effect on *F* or *Ŝ*. The Vpli model has a substantial effect on all the statistics. These results are representative of those from simulations with other values of θ.

**Figure 7.**
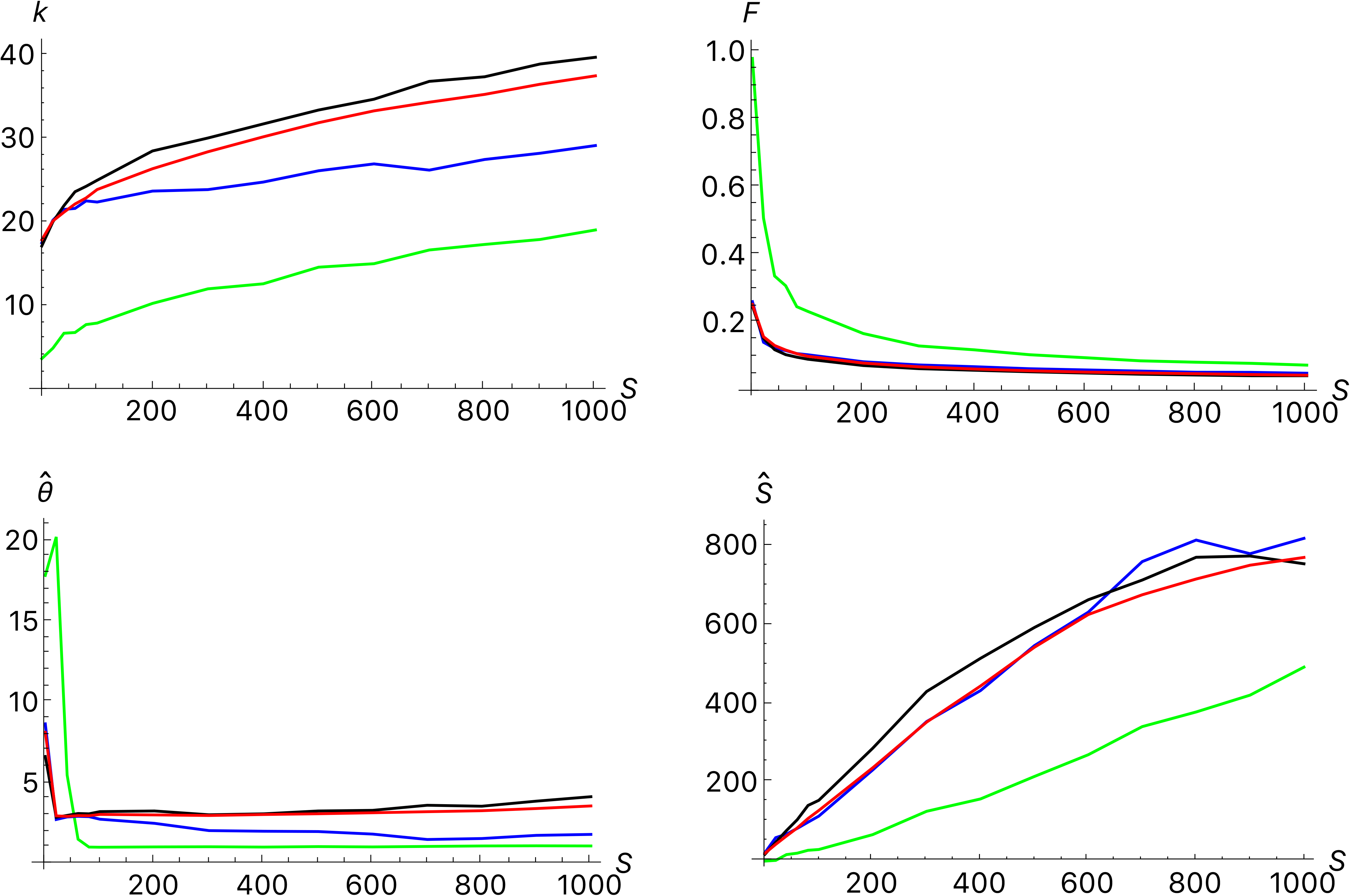
Comparison of models of asymmetric balancing selection. In all cases θ=3. Each model was run for a 100,000 generation burn-in period and then the simulation was run for an additional 1,000,000 generations. Samples of 500 individuals were collected every 10,000 generations after the burn-in period beginning with generation 0. Each point shown is the average of the results for 101 samples. The curves in red show the results for the symmetric model. The green curves show the results for the variable pleiotropy model (Vpli) with *r_max_* = 0.05. The blue curves show the results for the variable heterozygosity model (Vhet) with *u_max_* = 1. The black curves show the results for the variable homozygosity model (Vhom) with *v_max_* = 1.

We can understand these results better by plotting the allele frequencies against values of the variable parameter in each model, *v_i_*, *u_i_* or *r_i_*. Figure 8 shows snapshots of the simulated populations for samples of 500 individuals taken 400,000 generations after the end of burn-in period. For the Vhom model, there is a slight tendency for alleles with smaller *v_i_* to be in higher frequency, especially for stronger selection. For the Vhet and Vpli models, there is a strong relationship between allelic effect and allele frequency. For the Vhet model, alleles with large *u_i_* are in higher frequency while, for the Vpli model, alleles with small *r_i_* are in higher frequency.

**Figure 8.**
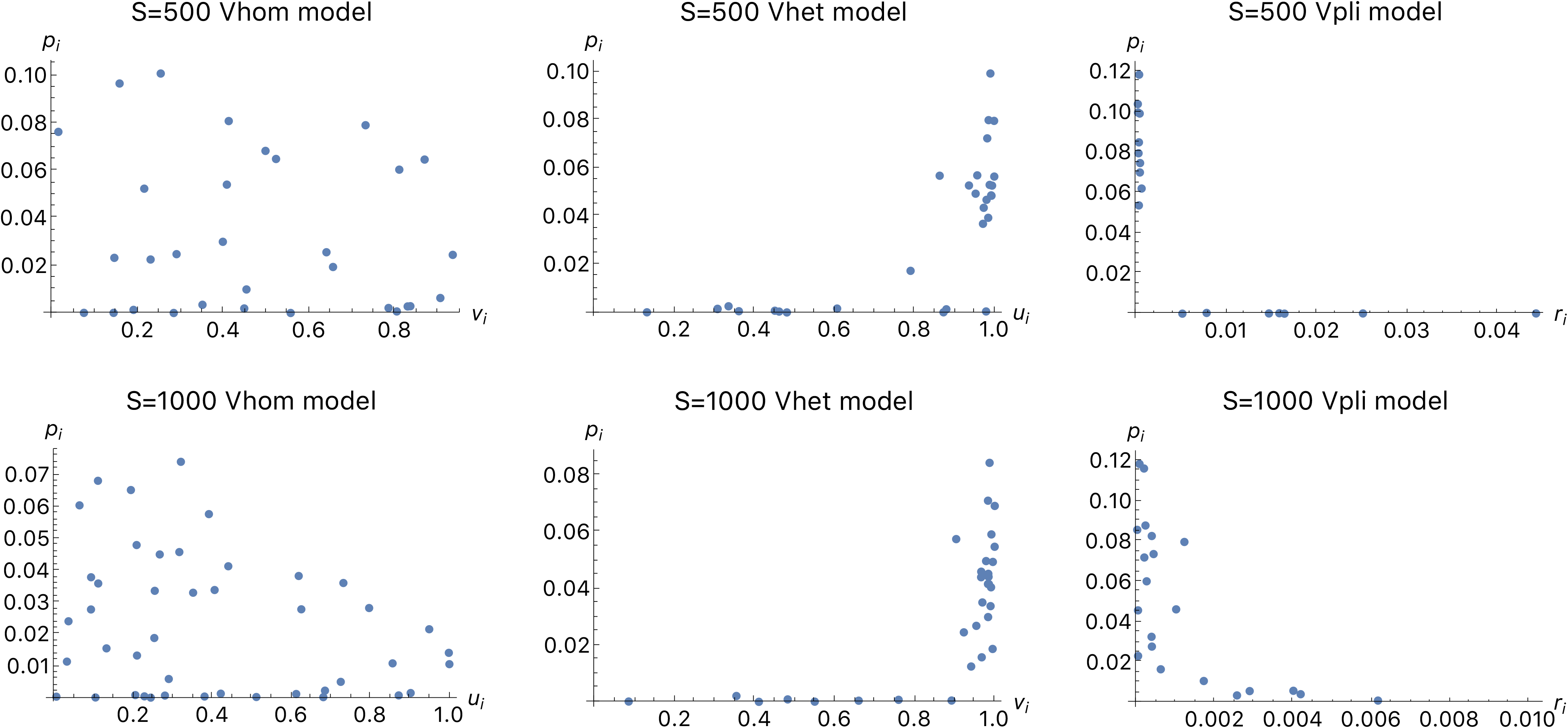
Allele frequency plotted against variable selection parameters for three models of asymmetric selection. A sample of 500 individuals was taken 400,000 generations after the end of the burn-in period in each replicate simulation of the variable homozygosity (Vhom), variable heterozygosity (Vhet) and variable pleiotropy (Vpli) models described in the text. In all cases θ=3. In the Vhom model *v_max_* = 1; in the Vhet model *u_max_* = 1; and in the Vpli model *r_max_* = 0.05. Note the different scales used for the horizontal axis of the Vpli model results.

These patterns are found because of the tendency for selection to increase mean fitness in a single-locus model when relative fitnesses are constant, a general principle in population genetics (Kingman 1961). In all three models, alleles differ in their effect on fitness when they arise by mutation. But those alleles that tend to increase mean fitness by more will be retained longer than those that do not. The result is a balance between the introduction of alleles by mutation and their stochastic loss at a rate that depends on their effect on mean fitness.

The difference between the Vhom model and the other two can be explained by the relatively weak dependence of 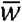 on the *v_i_*. In Equation (5), 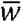 is reduced by 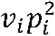 for allele *A_i_*. Because allele frequencies in these models tend to be small, the effect of each allele on mean fitness in the Vhom model is quite small. In contrast, the increase in 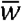 in the Vhet model for allele *A_i_* is *p_i_*(1 – *p_i_*)*u_i_* and hence is proportional to *p_i_* instead of 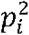. Although less obvious in Equation (7), the dependence of 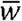 on *r_i_* is also roughly proportional to *p_i_* in the Vpli model. These patterns are consistent with the intuition that variation in allelic effects are least important in the Vhom model because homozygous individuals are relatively infrequent. Variation in allelic effects is more important in the Vhet model because all heterozygous individuals are affected and they are much more frequent. The effect is larger still in the Vpli model because both homozygotes and heterozygotes are affected.

Although there is a tendency to increase 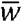 in these models, the maximum is never reached. Alleles that contribute most to increasing 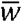 will still be lost because of genetic drift. A balance is achieved between the introduction of alleles by mutation and their eventual stochastic loss. One way to characterize this balance is with the weighted average of the random parameter: 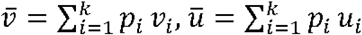, and 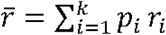. In simulations with no burn-in period, these quantities stabilize by roughly 20,000 generations, as shown in Figure 9. There is no tendency to increase after that time.

**Figure 9.**
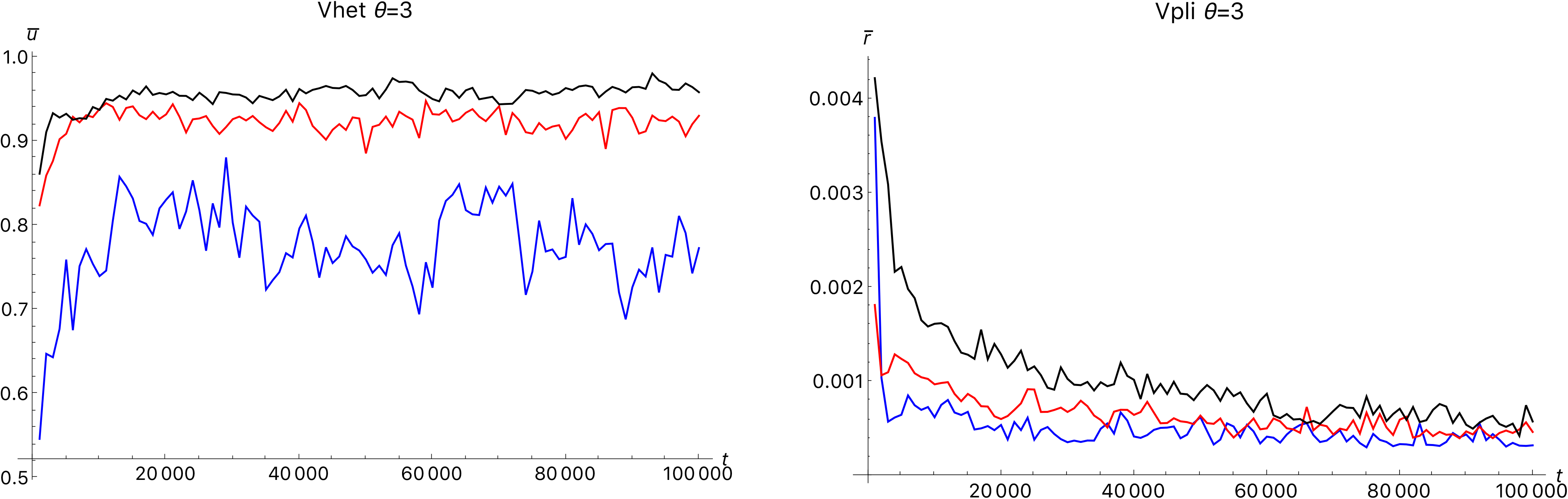
Weighted average values of *u_i_* and *r_i_* in simulations of the Vhet (with *u_max_* = 1) and Vpli (with *r_max_* = 0.05) models with no burn-in period. In both parts, θ=3 and *N*=10,000. Initially, each replicate was fixed for a single allele. Results for *S*=1000 (black), *S*=500 (red) and *S*=100 (blue) are shown.

An additional result from simulations of the Vhet and Vpli models is that they can both lead to more long-lasting alleles than do comparable symmetric models for moderate mutation rates, θ=0.5 or larger. For θ=0.1, the reverse is true. Figure 10 compares histograms of allele ages with and without variation in allelic fitness. For the Vhet model with *u_i_* distributed uniformly on (0,1), the appropriate model for comparison is one in which *u_i_* = 1 because *u_i_* is nearly 1 for alleles in moderate frequency, as shown in Figure 8. For the Vpli model, the model comparable to one in which *r_i_* is uniform on (0,0.05) is one in which *r_i_*=0. Table 2 quantifies the general patterns found. In Figure 10, there are fewer alleles overall when the selection coefficients are random because the effective mutation rate is smaller. Some mutations reduce mean fitness sufficiently that they are quickly lost. Nevertheless, some of the surviving mutations persist for longer times than in the models with constant selection coefficients.

**Figure 10.**
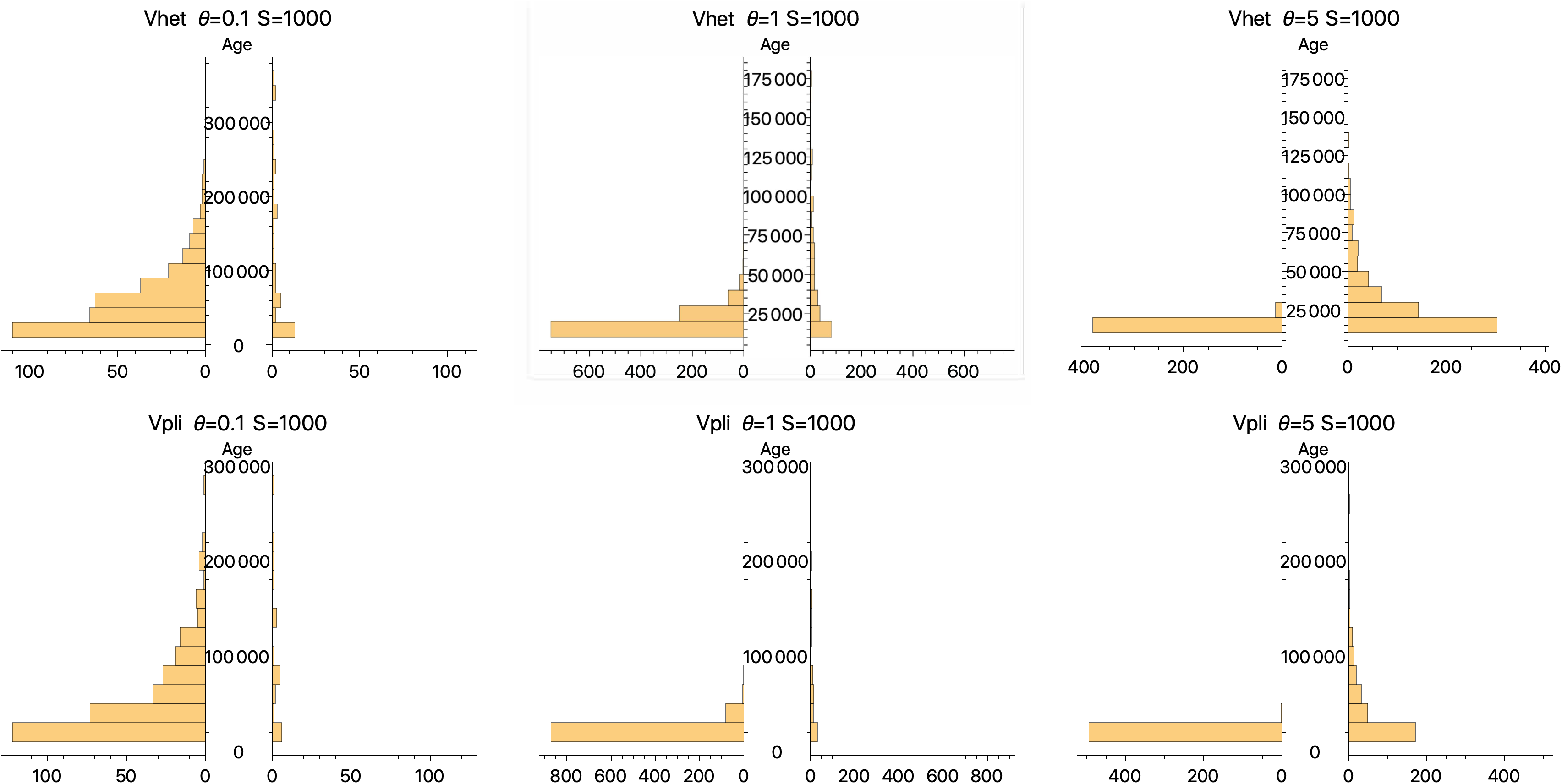
Paired histograms of alleles whose ages exceeded 100,000 generations. In all cases, a population of 10,000 individuals was simulated for 1,000,000 generations after a 100,000 generation burn-in period. The ages for all alleles that arose by mutation after the end of the burn-in period and were lost before the end of the 1,000,000 generations were recorded. In each of the paired histograms, the ones on the left are for *u_i_* = 1 (Vhet) or *r_i_* = 0 (Vpli) and the ones on the right are for randomly generated values. For the Vhet model, *u_i_* was uniformly distributed on (0,1) and for the Vpli model *r_i_* was uniformly distributed on (0,05).

## Trans-species polymorphism

Models of balancing selection can easily be made to generate high heterozygosity and large numbers of alleles by increasing the mutation rate and selection intensity. However, a relatively high mutation rate even when combined with very strong balancing selection results in a high turnover of alleles. Many alleles are maintained at any time but they do not stay in the population long enough to account for trans-specific polymorphism (TSP). For humans and chimps to share alleles, those alleles had to have persisted in both populations for six or more million years, more than 300,000 generations assuming an average generation time of 20 years.

Population geneticists have been aware of this problem. (Radwan *et al*. 2020) One solution is to assume that parameter values lie within a restricted range for which both a large number of alleles and long allelic persistence times are possible. Takahata (1990) used his analytic theory to show that low mutation rates and very strong selection together are needed. He estimated that the mutation rate in the peptide binding regions of HLA loci to be roughly 10^-7^ per generation implying *θ* = 0.004 if the effective population size is 10,000. With *s*=0.1 (*S*=2000), the expected time to loss of a selected allele is roughly 600,000 generations, more than enough to account for TSP between humans and chimps. (Takahata 1990) For those parameter values, the effective number of allele (*n_e_*=1/*F*) exceeds 20, which is also consistent with observatons.

Although Takahata’s results can account for TSP, they can do so only for a restricted range of parameter values: θ must be very small, much less than 1, and *s* must be very large (0.01 or greater) and that has to be true for every MHC locus that exhibits TSP. Furthermore, the parameter values that predict TSP also predict that allele frequencies are quite evenly distributed. We can quantify the evenness of the distribution of allele frequencies by computing the product *kF*. If frequencies are equal, this product will be one because *p_i_* = 1/*k* and hence *F*=1/*k*. This product exceeds 1 by an amount that reflects the unevenness of the allele frequencies. Figure 11 shows some values from the simulations presented earlier. If θ=0.1, *kF* approaches 1 for even moderate selection intensities. For larger values of θ, *kF* remains substantially larger than 1 even for *S*=1000. In the 1000 Genomes data, Table 1 shows that observed values of *kF* are greater than 1 and often greater than 2.

**Figure 11.**
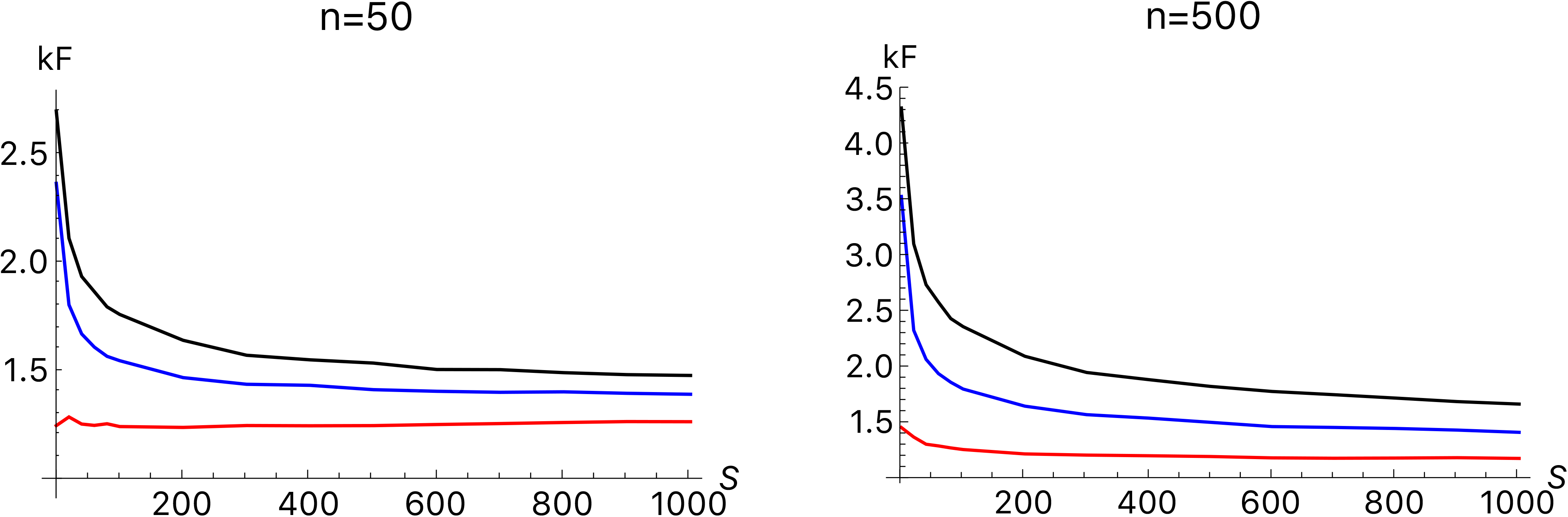
Values of the product of *k* (the number of alleles) and *F* (the homozygosity) for the symmetric model of balancing selection. The results are obtained from the same simulations as shown in Figures 1-3. Results are shown for three different mutation rates, θ=0.1 (red), 1 (blue) and 3 (black).

The simulation results presented above suggest another possible explanation for TSP, variation among alleles in their contribution to fitness. For the Vhet and Vpli models, variation among alleles greatly increases the chance that some alleles will remain in the population for a long time, much longer than is found for comparable symmetric models (Figure 10 and Table 2).

## Discussion and Conclusion

This paper generalizes the Kimura and Crow (1964) symmetric infinite-alleles model of balancing selection in an equilibrium population. First, it uses the analytic results of Muirhead and Wakeley (2009) as the basis of a composite likelihood method for jointly estimating the scaled mutation rate (θ = 4*N*μ) and the scaled selection coefficient (*S* = 2*Ns*). The method applied to data simulated under the symmetric equilibrium model results in estimates that are on average close to the true values. There is considerable variation of the estimates about the true values however. Roughly speaking, estimates of *θ* are within a factor of 2 of the true value while estimates of *S* are only within a factor of five or ten of the true value. There is limited information in the allele frequency spectrum in a population sampled at a single time.

The application of this method to HLA data presented by Gourraud et al. (2014) yielded a range of parameter estimates. As is predicted by the simulation results, the range of variation across populations in 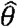 is smaller than the range of variation in *Ŝ* for each locus. There is some consistency in 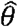 within loci. For example, 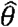 values for HLA-DQA1 are substantially smaller than for HLA-B. There is less consistency in values of *Ŝ* for each locus, but values for HLA-B tend to be larger than for other loci. We can tentatively conclude that the variability at HLA-B reflects stronger selection than at the other loci examined rather than a higher mutation rate.

There is a weak relationship between *Ŝ* and the outcome of the Ewens-Watterson test of neutrality. Samples for which *Ŝ* exceeds 100 generally reject neutrality at the 5% level if 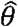 is small. For larger 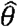, some cases with large *Ŝ* do not reject neutrality. That pattern is also predicted by the simulations. They showed that higher mutation rates reduce the power of the Ewens-Watterson test for a given *S*. The failure to reject neutrality in a sample does not necessarily indicate that selection is weak, only that the Ewens-Watterson test has relatively low statistical power in some cases.

The simulations of models with population growth and population subdivision showed that 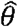 and *Ŝ* are affected by deviations from the equilibrium model. Even in these relatively simple demographic models, the estimates do not reflect the local effective population size at the time the samples were taken. There is no reason to think that more complex and realistic demographic assumptions would alter that conclusion. Instead, 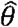 and *Ŝ* obtained from a single sample have to be interpreted as indicating that the allele frequency spectrum is comparable to one expected in a stable isolated population with those parameter values. When several loci from the same population are analyzed, the values of 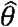 indicate the relative mutation rates because all loci experienced the same demographic history. For example, the results in Table 1 indicate that the mutation rate at HLA-B is probably five to ten times higher than the mutation rate at HLA-DQB1.

The estimates of θ and *S* in Table 1 differ from those obtained by Yasukochi and Satta (2013). They estimated θ (4*M* in their notation) from inferred rates of replacement substitutions (*K_B_*), the selective advantage of replacement to silent substitutions (*γ*), and the numbers of sites in the peptide binding regions of different HLA class I and II loci. Their estimates of θ ranged from 0.04 for HLA-B, HLA-DRB1 and DPB1 to 0.56 for HLA-DQB1 and 0.6 for HLA-C. These estimates are different from those shown in Table 1 which are largely reflecting the numbers of rare alleles. The methods are so different that it is difficult to know why the estimates of θ are different. The rate of gene conversion would not be accounted for in the estimate of *K_B_*. Even the relative rates are dissimilar. In Table 1, HLA-B has the largest estimates of θ while Yasukochi and Satta (2013) inferred that it had one of the smallest estimated rates. The reverse is true for HLA-DQB1.

Yasukochi and Satta’s (2013) estimates of *S* are larger than those in Table 1. They range from roughly 500 for HLA-DQB1 to well over 1000 for the other loci. The difference in those estimates are probably attributable to the fact that Yasukochi and Satta’s estimates of θ are smaller than those in Table 1. Yasukochi and Satta assumed that Takahata’s (1990) analytic approximation was valid, which it would be if the mutation rates are as small as their estimates. One thing that argues against estimates of θ and *S* obtained by Yasukochi and Satta is that, if accurate, the Ewens-Watterson test should always reject the hypothesis of neutrality. Both here (Table 1) and the previous analysis by Solberg et al. (2008), neutrality is sometimes rejected but often is not. Furthermore, values of the product *kF* shown in Table 1 are generally larger than is expected when selection is strong and mutation is weak. Both my analysis and that of Yasukochi and Satta’s (2013) reflect different aspects of the data and hence are not easily compared. Further analysis is clearly needed to reconcile these differences.

The models of asymmetric selection analyzed here suggest that variation in allelic effects on fitness can have two consequences. First, differences in the allelic contribution to fitness result in a preferential retention of alleles that increase mean fitness. Consequently, asymmetry in the mutation process does not necessarily lead to substantial deviations from the predictions of a symmetric model. Here, the asymmetry is imposed at the mutation stage, but more realistic models that show how such asymmetry can arise have been considered. For example, De Boer et al. (2004) assume allelic fitness depends on the relationship between the sequence of the peptide binding region and the spectrum of antigen sequences. In fact, Figure 2 of that paper shows results similar to those in Figure 8 above. Stefan et al. (2019) generalized the De Boer et al. model and emphasized the advantages of divergent alleles that would bind with a wider range of antigens. Stefan et al. (2019) also found that more fit alleles would tend to accumulate at higher frequencies. Lighten et al. (2017) make similar assumptions. Siljestam and Rueffler (2019) assume there is a tradeoff between an allele’s ability to confer resistance to specific pathogens and the number of pathogens that can be resisted. These models and similar ones represent hypotheses about how fitness differences among alleles arise and are maintained by pathogen interaction and coevolution. Once the alleles are present in the population and their fitnesses are determined by interactions with pathogens, alleles are still governed by standard population genetic processes.

One class of models not analyzed here are models of divergent-allele advantage (DAA) first proposed by Wakeland et al. (1990). There is empirical support for the DAA model. (Pierini and Lenz 2018) Some analyses of DAA models concluded that they do not augment variation or readily lead to trans-specific polymorphism. (Lau *et al*. 2015; Ejsmond *et al*. 2018) Uyenoyama (2003) reached a similar conclusion about the evolution of self-incompatibility alleles, which experience fertility selection comparable to very strong balancing selection (Vekemans and Slatkin 1994). Others analyses, including those of Lighten et al. (2017) and Stefan et al. (2019), reach the opposite conclusion that a DAA can greatly increase allelic diversity. These different conclusions can be reconciled by considering the effect of DAA on mean fitness. A simple generalization of the Vhet model that allows for DAA is one in which the relative fitness of a homozygote is 1 – *s* (as in the symmetric model) and the relative fitness of an *A_i_A_j_* individual has relative fitness 1 + *su_ij_*, where the *u_ij_* are chosen to allow for arbitrary interactions among alleles. In the DAA model the *u_ij_* would be smaller for some pairs of alleles to reflect the fact that they are more similar each other or are more recently diverged. Values of the *u_ij_* for other pairs of alleles would be larger because they are more dissimilar and can bind to a wider range of antigens. For this model, the average relative fitness is 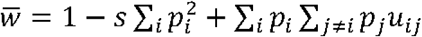, which is an increasing function of the *u_ij_*. (cf, Equation 6). Alleles that tend to increase 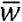 will be preferentially retained. That is what is seen in the simulation results of Stefan et al. (2019).

Models of coevolution with pathogens are somewhat different from the asymmetric models analyzed here because each allele’s contribution to fitness changes with time as pathogens coevolve in response to the presence of new alleles in relatively high frequency. Ejsmond and Radwan (2015) noted that coevolutionary models are similar to the Red-Queen process in evolutionary biology because MHC alleles continually need to keep up with new challenges presented by coevolving pathogens.

The results from the analysis of asymmetric models have implications for trans-specific polymorphisms. The three models examined here are not intended to be realistic. Instead they are intended to explore the consequences of variation in allelic contributions to fitness. One of those consequences is that long term persistence of a few alleles is possible even with very small differences in allelic fitness. Stochastic loss of an allele is a sensitive function of selection intensity (Takahata 1990) so slight differences in allelic contributions to fitness affect persistence times disproportionately. Any biological constraint on allelic fitness arising from their role in pathogen defense could maintain larger differences among alleles and hence lead to much larger differences in persistence times. Trans-specific polymorphisms are then possible under a wider range of parameter values than is suggested by the symmetric equilibrium model.

Although I have tried to relax the assumptions of the standard model of balancing selection in several ways, the models explored in this paper are still quite simple. They are intended to indicate the ways in which alterations to the symmetric model affect predictions about the maintenance of variation in MHC loci in humans and other vertebrates.

## Program availability

Copies of the C programs used for the simulations and data analysis will be made freely available at GitHub.com.

## Acknowledgements

I thank J. Felsenstein, S. J. Mack and C. Poux for helpful comments on an earlier version of this paper, and H. Erlich, M. Fernandez-Vina, J. Hollenbeck, M. Maiers, K. Osoegawa, R. Single, and G. J. Thomson for helpful discussions during the early stages of the work presented here.

